# SiaRNA: A Siamese Neural Network with Bidirectional Cross-Attention for Pairwise siRNA-mRNA Efficacy Prediction

**DOI:** 10.64898/2025.12.13.694094

**Authors:** Sapireddy Vaishnavi, Nathi Rajkumar, Meruva Venkata Harshit, Nannapuraju Varun Raju, Basangari Bhargava Chary, Kondaparthi Vani

**Author notes:** Authors contributed equally (Sapireddy Vaishnavi, Nathi Rajkumar, Meruva Venkata Harshit, Nannapuraju Varun Raju).

## Abstract

Small interfering RNA (siRNA) therapeutics have extraordinary potential for targeted gene silencing. They mediate post-transcriptional gene regulation by binding to complementary messenger RNA (mRNA) sequences and degrading them, thereby preventing the production of unwanted proteins. Recent machine learning and deep learning frameworks for predicting siRNA efficacy have only achieved moderate success as these models solely rely either on handcrafted features or on sequential relations and therefore cannot capture the full complexity of siRNA-mRNA interactions. In this context, we propose SiaRNA, which uses a Siamese Neural Network for feature-derived representations and a bidirectional cross-attention mechanism for sequence-level relationships. It uniquely identifies mRNAs and their corresponding siRNAs as paired entities, allowing unified and context-aware modeling. Unlike previous models, which discard 2-nucleotide (2-nt) overhangs at the 3’ end while using 21-nt efficacy labels, SiaRNA both trains and tests on 21-nt sequences to ensure biologically consistent predictions. Our model sets a new performance benchmark, outperforming previous state-of-the-art models. SiaRNA is trained on the HUVK dataset achieving an accuracy of 0.881, while its generalization has been confirmed by testing on the independent Simone dataset. These results prove SiaRNA’s potential as a reliable and biologically accurate framework to guide siRNA design and improve therapeutic outcomes.

## 1. Introduction

RNA interference (RNAi) [1] is a natural cellular defense mechanism found across eukaryotic organisms that regulates gene expression at post-transcriptional level. Transcription is the first step in gene expression, where genetic information encoded in DNA is copied into messenger RNA (mRNA) within the cell nucleus by RNA polymerase enzymes. The newly synthesized mRNA molecule then exits the nucleus and enters the cytoplasm, where translation occurs, during which the ribosomes decode the mRNA to synthesize the corresponding proteins.

Proteins are essential molecules in the body, responsible for carrying out almost every cellular function. However, protein production can be disrupted due to genetic mutations, transcription or translation errors [2] and viral infections [3], leading to serious conditions such as cancer [4], genetic disorder, metabolic disorder and neurodegenerative diseases. To control such abnormalities, cells employ the RNA interference (RNAi) pathway, which regulates protein production through double-stranded RNA (dsRNA). dsRNA is a type of RNA that can naturally form within the cells, such as during viral infections. Inside the cell, the long double-stranded RNA (dsRNA) is recognized by an enzyme called Dicer, a special type of ribonuclease (RNase). Dicer cleaves the dsRNA into smaller fragments known as small interfering RNAs (siRNAs)[5], typically 19-23 nucleotides in length. To induce RNAi, small interfering RNAs (siRNAs) or short hairpin RNAs (shRNAs) are introduced into the cell, where they mimic the natural products of the RNAi pathway. These siRNAs are then incorporated into the RNA-induced silencing complex (RISC), which is a multi-protein effector complex that carries out gene silencing. Inside RISC, the siRNA loses one of its strands, called passenger strand, while the other strand, guide strand, is retained. The guide strand serves as a template to find complementary mRNA. By means of base-pairing interactions, it scans the cellular mRNA to identify target sequences. Once the siRNA-mRNA duplex forms, Argonaute 2 (AGO2) protein, which is the catalytic core of RISC, cleaves the target mRNA at the site of the complementary region. The resulting mRNA fragments are then degraded by RNA-degrading enzymes that prevent the translation of mRNA into protein, thus silencing the target gene. This selective process makes sure that only specific genes are silenced, allowing the cell to accurately control the synthesis of proteins and maintain cellular balance.

Advancements in computational biology and machine learning have helped grow the study of RNA interference beyond traditional experimental methods. Various state-of-the-art computational models have been developed for the prediction and analysis of siRNA-mRNA interactions like s-Biopredsi [6], DSIR [7] and i-Score [8]. These developed models for siRNA efficacy prediction span rule-based frameworks to neural network architectures and advanced data mining techniques. Despite the progress made with the help of these frameworks, there are still many persistent challenges such as inconsistency in performance across datasets due to variations in experimental procedures, high dependency on feature selection and an inability to model non-linear interactions.

To address these limitations, Convolution Neural Network (CNN) [9] based models were employed to analyze sequence context along with thermodynamic properties which provided better results than previous models but still struggled in capturing high level interaction. Later, Graph Neural Networks (GNN) became prominent in this field due to their ability in capturing intricate relationships between multiple siRNA-mRNA pairs through message passing. Models such as GNN4siRNA [10] and siRNADiscovery [11] have since become state-of-the-art GNN models. GNNs require large and diverse data which is difficult to obtain in this domain and cannot be efficiently parallelized due to complex dependencies inherent in graph structures. Moreover, graph-based models tend to lose key structural details when handling imbalanced data, leading to over-smoothing [12] or signal dilution, resulting in degraded performance.

To overcome these challenges, we introduce SiaRNA, a deep learning model that integrates a Siamese Neural Network (SNN) [13] with bi-directional cross-attention [14] mechanism. SiaRNA effectively captures both pair-wise dependencies and sequential relations. In addition, thermodynamic features are integrated before the final prediction stage to the outputs of the SNN and bi-directional cross attention modules. The model has been trained and evaluated exclusively on 21-nucleotide (21-nt) siRNAs, consistent with experimentally validated standards.

## 2. Results and Discussion

### 2.1. Model Training

All experiments were conducted on a NVIDIA A100 GPU, which provided the computational capacity necessary to train the model on high-dimensional biological sequences. From the total 2816 siRNA-mRNA pairs in Dataset_HUVK, 1971 were used for training, 423 for validation and 422 for testing.

The model was trained using the Adam optimizer, chosen for its effective convergence. Gradient clipping was used to ensure numeric stability while training. This technique mitigates exploding gradients which is a common source of instability in deep networks. In addition, layer normalization was used to stabilize activation across layers which improves training stability. This mechanism guarantees that the model’s weights are updated smoothly leading to more stable convergence.

### 2.2. Hyperparameter configuration

The model’s training was controlled by a set of precise hyperparameters. These parameters define the dimensions of all layers, from the initial high-dimensional projection of input features to the output dimensions of the shared Siamese embedder. They also specify the number of heads used in the cross-attention module and the layer sizes within the fusion layer and the regression Multi-Layer Perceptron (MLP).

**Table 1.** Summary of Model Hyperparameters. Table 1 lists the key training parameters and model parameters along with their corresponding values used during model development.

| Category | Hyperparameter | Value |
| --- | --- | --- |
| Data & Features | Max siRNA Length | 21 |
|  | Max mRNA Length | 9756 |
|  | mRNA Slice Length | 57 |
| Model Architecture | SNN Projection Dimension | 256 |
|  | Hidden Dimension | 64 |
|  | Embedding Dimension | 32 |
|  | Attention Heads | 4 |
|  | Fusion Dimension | 128 |
|  | MLP Dimension | 64 |
| Training | Learning Rate | 1e-3 |
|  | Epochs | 20 |
|  | Batch Size | 32 |
|  | Gradient Clip Norm | 1.0 |
| Loss Function | Lambda Metric | 0.1 |
|  | Contrastive Margin | 1.0 |

### 2.3. Performance on Dataset_HUVK

To test the model, we benchmarked its performance on Dataset_HUVK against siRNADiscovery, using 422 siRNA-mRNA pairs (15% of the total dataset). SiaRNA yielded an average MSE of 0.021, PCC of 0.769, SPCC of 0.775 and AUC of 0.881. Meanwhile, siRNADiscovery achieved an average MSE of 0.020, PCC of 0.770, SPCC of 0.771 and AUC of 0.874. The results demonstrate that SiaRNA achieves performance that is highly competitive and on par with the current benchmarks. Among the compared models, i-Score exhibited the weakest performance, having PCC, SPCC and MSE of 0.602, 0.623 and 0.064 respectively.

**Table 2.**
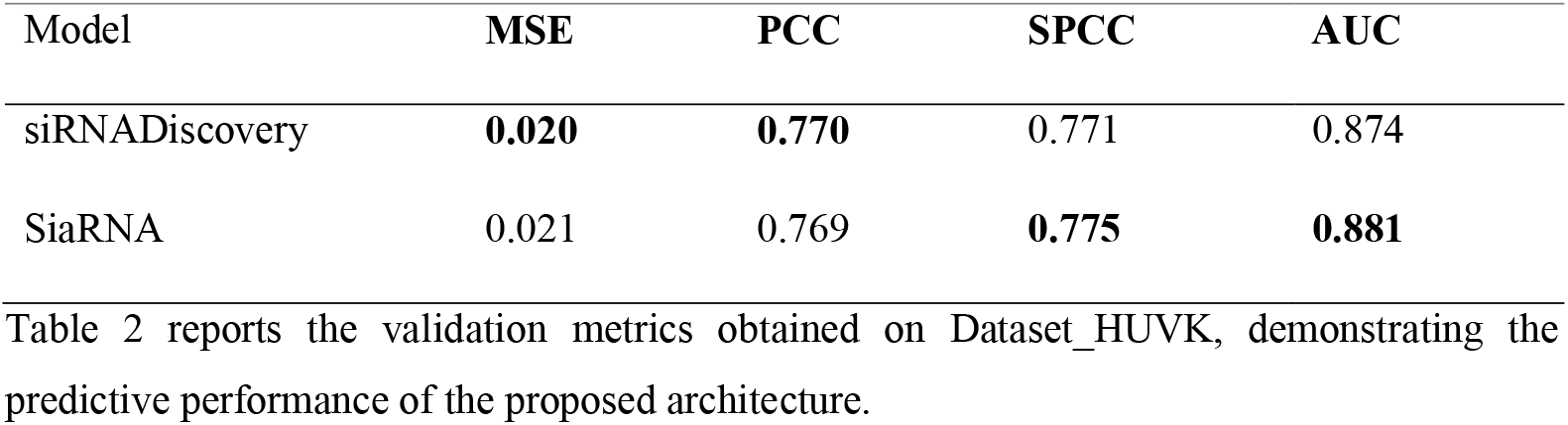
Validation results on Dataset_HUVK.

| Model | MSE | PCC | SPCC | AUC |
| --- | --- | --- | --- | --- |
| siRNADiscovery | **0.020** | **0.770** | 0.771 | 0.874 |
| SiaRNA | 0.021 | 0.769 | **0.775** | **0.881** |

Table 2 reports the validation metrics obtained on Dataset\_HUVK, demonstrating the predictive performance of the proposed architecture.

### 2.4. Performance on Simone Dataset

To further assess the robustness and generalization of SiaRNA, it was evaluated on the independent external Simone dataset, which contained 322 siRNA-mRNA pairs. The results show that SiaRNA demonstrates strong predictive capability and achieves consistently higher performance across the evaluation metrics. SiaRNA achieves a PCC of 0.5163, SPCC of 0.3849 and AUC of 0.7586, compared to siRNADiscovery’s PCC of 0.464, SPCC of 0.32 and AUC of 0.702. As seen in Figure 1, SiaRNA outperforms other state-of-the-art models on the Simone dataset. These findings suggest that SiaRNA provides improved generalization on unseen data and effectively addresses some of the limitations observed in previous architectures.

**Figure 1.**
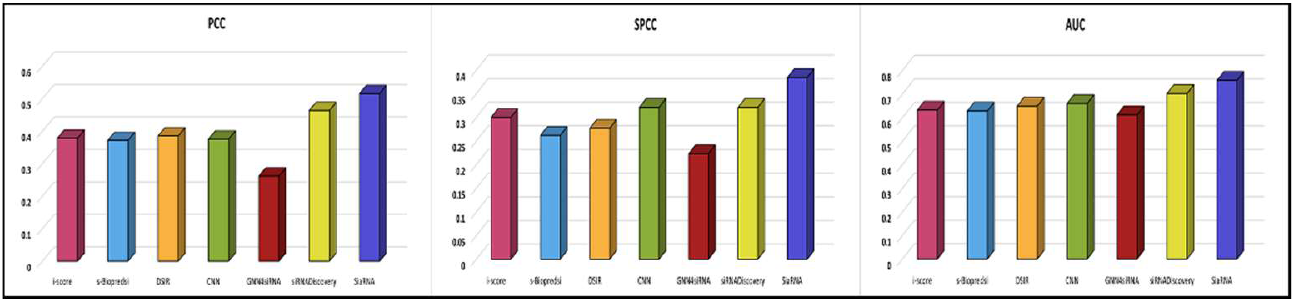
Comparative performance of models on Simone dataset.

Figure 1 represents three graphs comparing the performance of models against three important evaluation metrics, namely Pearson correlation coefficient (PCC), Area under the ROC curve (AUC), and Spearman correlation coefficient (SPCC). For each graph, the x-axis represents the models, while its y-axis represents the corresponding evaluation metric score. All three graphs compare SiaRNA to existing state-of-the-art models and demonstrate overall improvements in predictive accuracy and generalization on the external Simone dataset.

### 2.5. Ablation Study

An ablation study was conducted as a means to investigate the individual contribution of key model components separately. The key components considered were the bidirectional cross attentional module, thermodynamic features and contrastive loss. The ablation of these model components was chosen because they represent the conceptually and functionally distinct elements in the architecture. The observed results of the ablation study are visualized in Figure 2.

**Figure 2.**
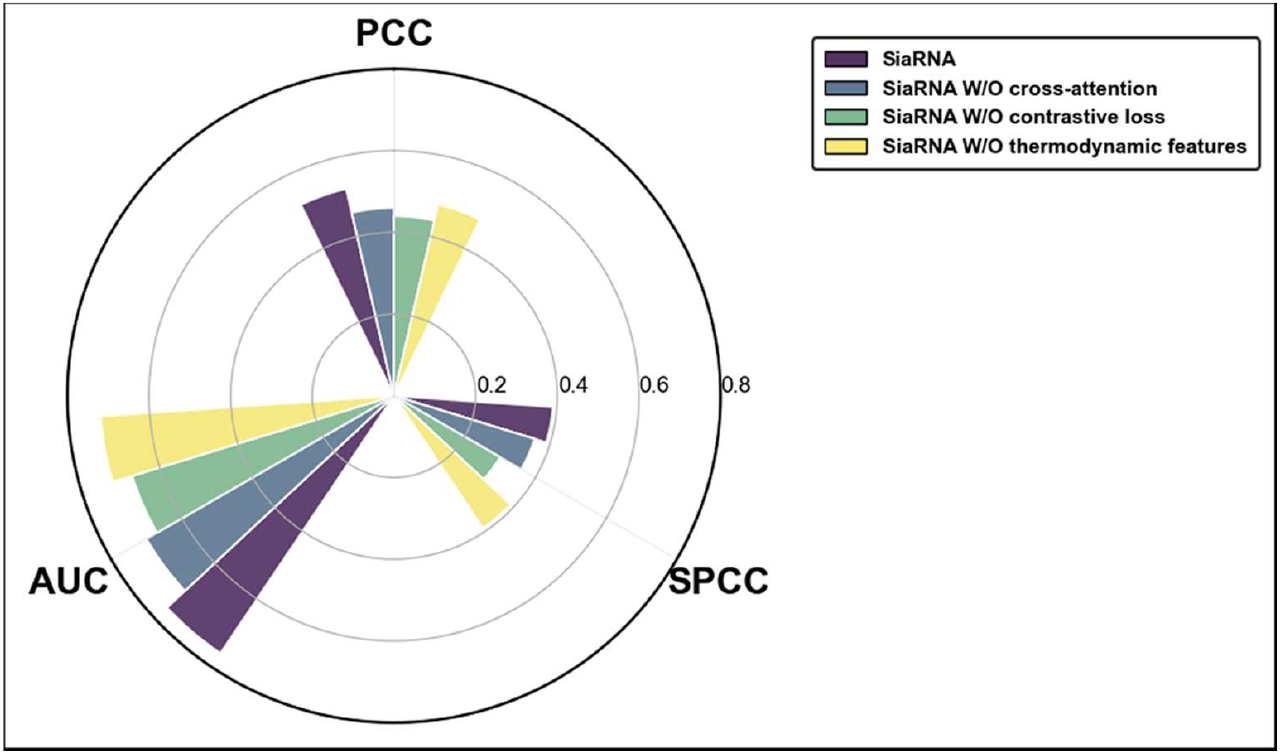
Comparison of Ablated variants.

The bidirectional cross-attention module is a core structural component that models the contextual dependencies between the mRNA and siRNA sequences. Evaluating its removal helps determine how much of the model’s predictive power comes from explicit sequence-level interaction learning. The thermodynamic feature module acts as an independent source of information derived from biophysical principles. By removing it, we investigate the degree to which the thermodynamic properties of the siRNA-mRNA pairs complement the learned features from the SNN and bidirectional cross-attention modules. Finally, the contrastive loss was ablated because it represents a novel addition to the learning objective, intended to enforce alignment between the siRNA and mRNA embeddings in the shared latent space. The removal of this component shows precisely its contribution in promoting generalization.

Figure 2 shows ablation experiments done to validate the contribution of key modules and components of the proposed model architecture. In this, “W/O” stands for “without”, implying that certain modules, such as cross-attention, thermodynamic features, or contrastive loss, were removed to test their contribution individually.

## 3. Methodology

### 3.1. Overview

SiaRNA is designed to capture thermodynamic, compositional, and contextual dependencies between siRNA and its target mRNA. Specifically, the model first processes a set of handcrafted input features using a Siamese Neural Network. In parallel, the raw nucleotide sequences of siRNA and the corresponding mRNA region are numerically encoded based on nucleotide identity and passed through a shared bi-directional cross attention mechanism. The outputs from the feature-based and attention-based modules are then fused with thermodynamic features [15] and passed through fully connected layers to predict the final efficacy. The hybrid design enables the model to leverage both biologically interpretable features and to learn deep representations, achieving a comprehensive understanding of siRNA-mRNA interactions.

### 3.2. Data Collection and Characteristics

To train the model, 2816 unique siRNA-mRNA pairs and their respective efficacies from Massimo *et al [10]* are used, which originally come from the studies of Huesken [6], Harborth [16], Ui-Tei [17], Vickers [18], and Khovorova [19]. This combined dataset is referred to as Dataset_HUVK. Further, the Simone dataset [20], consisting of 322 unique siRNA-mRNA pairs is used to test the model.

Both the training and testing datasets being used have siRNAs of length 21-nt only. Earlier computational approaches often relied on truncated 19-nt sequences for convenience and compatibility with older datasets, a practice that introduces inconsistencies when efficacy labels derived from 21-nt experiments are applied to 19-nt input sequences [21]. Dataset_HUVK and Simone dataset, which are being employed in the current study provide efficacy measurements for 21-nt siRNAs which include the characteristic two nucleotide overhangs at the 3’ end, essential for recognition and loading of the siRNA by the RNA-induced silencing complex (RISC). Experimental evidence suggests that 21-nt siRNAs exhibit superior stability and silencing efficacy compared to their 19-nt counterparts, as the 3’ overhang facilitates AgO2-mediated strand loading and target recognition [22]. Therefore, using the full 21-nt representation ensures biological consistency between the experimental conditions and model inputs, improving both predictive validity and real-world applicability.

A 10-fold cross-validation strategy is employed, splitting Dataset_HUVK (2816 siRNA-mRNA pairs) in a 70:15:15 ratio into training (1971 siRNA-mRNA pairs), validation (423 siRNA-mRNA pairs) and test (422 siRNA-mRNA pairs) splits. The data is split ten times and each time using different random seeds to generate ten distinct folds of data which reduces sampling bias and ensures reliable evaluation of performance.

### 3.3. Feature Engineering

The proposed framework utilizes two distinct input streams - one for the feature-engineered representations through the Siamese Neural Network (SNN) block and another for sequence-based representations for the Bidirectional Cross-Attention block.

#### 3.3.1. Feature Inputs for Siamese Neural Network

For the SNN component, comprehensive feature vectors are constructed for siRNA and mRNA using their sequences.

##### Sequence Composition and Encoding

- One-Hot Encoding: To capture positional nucleotide identity, both the siRNA and mRNA sequences are encoded using binary one-hot encoding scheme with each base (N, A, G, C, U/T) being assigned a unique four-dimensional vector: N=<0,0,0,0>, A=<1,0,0,0>, C=<0,1,0,0>, G=<0,0,1,0>, U/T=<0,0,0,1>(PMID: 27307608).
- Nucleotide Frequencies (siRNA Only): To quantify the influence of short sequence motifs on siRNA function, the frequency of short segments (k-mers) is calculated for the sequences. Previous work shows that k-mers significantly influence knock-down efficacy [23]. For example, dimer and trimer compositions have emerged as strong predictors of potency in machine learning models [24] and specific motifs such as UCC or A/U-rich regions - are statistically associated with enhanced silencing efficacy [25]. The calculated frequencies are: 1-mer (A,U,C,G) with 4 motifs, 2-mer (e.g., AU, CG, etc.) with 16 possible motifs, 3-mer (e.g., AUC, UCG, etc.) with 64 possible motifs, 4-mer (e.g., AUCG, UCGA, etc.) with 256 possible motifs and 5-mer (e.g. AUCGA, UCGUA, etc.) with 1024 possible motifs
- G/C Percentages: GC content for both the siRNA guide strand and the mRNA target sequence is computed. GC content is a critical factor influencing the stability of binding between the siRNA and mRNA. Low GC content can lead to weak, non-specific binding, while excessively high GC content can hinder the essential unwinding of the siRNA duplex by the RISC complex, thus impeding efficacy [25, 26].

##### Interaction and Rule-Based Features

- RNA-AGO2 Interaction Score: The score represents a quantitative measure of the RNA-protein interaction (RPI) between the RNA (siRNA or mRNA) transcript and the Argonaute 2 (AgO2) protein [27]. The score is computed using ZHMolGraph [28], a framework designed to capture the structural and chemical interactions between RNAs and proteins. The computed score represents the RISC loading potential of siRNA guide strand, which is a key biological feature for predicting efficacy.
- Rule Codes (siRNA Only): To encode the established knowledge that certain nucleotides at specific positions enhance or impair siRNA function, simplified rule codes are applied to the siRNA guide strand, based on the findings of He *et al* [20]. The original preference scores (1 for enhancing, -1 for reducing, 0 for no preference) are converted into unique-three dimensional binary vectors: 1=<0,0,1>, 0=<0,1,0>, - 1=<1,0,0>.

#### Structural Features

- Base Pairing Probabilities: The nucleotide base pairing within the individual strands (siRNA and mRNA) drives the formation of their secondary structures. This folding is biologically relevant because, within the cellular environment, both RNA transcripts exist primarily as folded species, not as linear strands. Both canonical (A-U, C-G) and non-canonical (G-U) interactions are considered. The RNAfold tool from ViennaRNA [29] package is used to obtain the base-pairing probability matrix for the individual siRNA and mRNA sequences. Due to the size and sparsity of these base-pairing probability matrices, truncated Singular Value Decomposition (SVD) is applied for dimensionality reduction, yielding compact representations of six dimensions for siRNA and one hundred dimensions for mRNA, which are subsequently used as model features.

#### 3.3.2. Feature Inputs for Bidirectional Cross Attention

For the bi-directional cross attention module, siRNA and mRNA sequences are represented as tokenized nucleotide indices [30]. Each siRNA (21-nt) is fully encoded, while for the mRNA, a fixed 57-nt [31] slice centered around the siRNA binding site is extracted. The mRNA slice has 28-nt upstream and 29-nt downstream of the target binding site. If the extracted mRNA sequence is shorter than 57-nt, suitable padding is applied to ensure compatible input length. Each nucleotide within these sequences is then numerically encoded into integer indices using a predefined mapping as: adenine (A) to 0, thymine (T) and uracil (U) to 1, guanine (G) to 2, cytosine (C) to 3 and ambiguous bases (N) to 4. This encoding is implemented by converting each nucleotide into its corresponding integer index and forming a tensor, which acts as the input to the attention mechanism. This representation allows the siRNA and target mRNA to be in uniform and computationally efficient format to analyze the contextual dependencies between them.

### 3.4. Architectural Framework and Core Components

The proposed architecture of SiaRNA integrates sequence-derived, contextual and thermodynamic information in order to achieve accurate modelling of siRNA-mRNA interactions. As shown in Figure 3, it consists of three key components: a Siamese Neural Network (SNN), a bidirectional cross-attention module and a thermodynamic-MLP component, where precomputed thermodynamic stability features are concatenated with the learned embeddings and processed through a MLP for final efficacy prediction.

**Figure 3.**
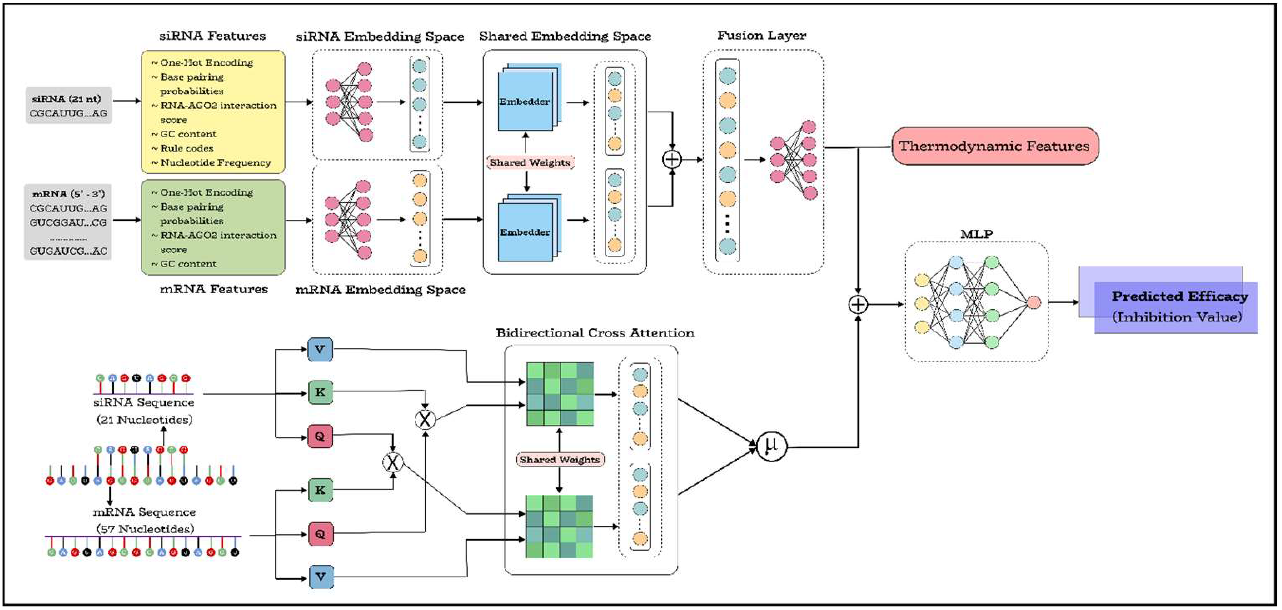
Architecture Diagram of SiaRNA.

Graph-based RNA interference models represent mRNAs as central hubs connected to multiple siRNAs, forming star-like topologies. While this approach captures network-level relationships, it may obscure pair-specific effects and limit inference on new siRNA-mRNA pairs. In contrast, the SiaRNA architecture treats each siRNA-mRNA interaction as an independent pair, allowing the model to learn specific binding patterns that arise from the joint sequence context [32] of the siRNA-mRNA pairs. This approach directly echoes the molecular mechanism of RNA interference (RNAi), where the functional outcome is determined by the compatibility between a single siRNA guide strand and its target mRNA sequence. Therefore, the model is better positioned to predict the efficacy of novel siRNA-mRNA pairs, which is crucial for practical siRNA design.

Figure 3 illustrates the end-to-end framework comprising two primary modules: a Siamese neural network for extracting pairwise characteristics and a bidirectional cross-attention module for capturing sequence-level relationships. The outputs from both modules are concatenated with thermodynamics features and passed through a multilayer perceptron (MLP) for final efficacy prediction.

#### 3.4.1. Siamese Neural Network

A Siamese Neural Network (SNN) is a specialized architecture that consists of two or more identical subnetworks, each sharing weights and parameters, designed to process pairs of inputs in parallel. While traditional neural networks predict specific outputs over individual inputs, SNNs are good at comparing features which makes them particularly effective for applications where the relationship or compatibility between two entities is to be studied.

The SNN architecture allows siRNA and mRNA to be encoded through parallel pathways before being projected into a shared latent representation. This design intuitively has a biological interpretation. For instance, when a given siRNA effectively silences its target, their shared embeddings will be closer together and in contrast for a weak pair, the shared embeddings will be farther away. This encourages biologically meaningful alignment between siRNA and mRNA features and enhances the model’s capacity to learn cross-molecular compatibility. This shared embedding mechanism thus enables the quantification of “functional proximity” between siRNA and mRNA sequences beyond surface-level similarity.

##### Linear Projection Layers

The linear projection layers provide the starting point for dealing with the raw handcrafted features of the siRNA and mRNA. These transform the features of different sizes and scales into fixed dimension vectors and project these vectors into the respective siRNA and mRNA embedding spaces. Independent processing of the handcrafted features allows the model to learn better by letting the siRNA and mRNA develop their own feature representations before interacting with each other.

##### Shared Embedder

The shared embedder component processes the standardized siRNA and mRNA features using the same set of weights for both inputs, projecting them into a shared embedding space, where the model is forced to learn a common language. This allows the model to measure the compatibility or “distance” between the siRNA and mRNA in the shared space, a step that is essential for learning patterns that define a successful interaction.

##### Fusion Layer

The Fusion Layer is designed to concatenate the shared representations of the siRNA and mRNA feature vectors into a single, cohesive vector. The combined vector is then fed into a fully connected layer which models the complex, non-linear interactions between the siRNA-mRNA pairs. The goal of this component is to move beyond the individual characteristics and create a unified representation (*h*_SNN_) that captures the relational properties of the siRNA-mRNA pairs.

#### 3.4.2. Bi-directional Cross Attention for Nucleotide-Level Interaction

The Bidirectional Cross-Attention mechanism is a very important part of the model, designed to simulate the subtle yet dynamic interaction between the 21-nt siRNA and its target 57-nt slice of the mRNA. The task of this component, beyond simple matching of sequences, is to learn complex position-dependent dependencies that influence binding affinity. This mechanism operates as a bi-directional, two-way process, effectively modelling the interaction between the two sequences to determine their compatibility.

The attention mechanism provides the model with the ability to selectively concentrate on the most relevant parts of the input. It computes a weighted sum of input features, where the weights indicate the importance of each input element for the current output. Mathematically, given queries *Q*, keys *K* and values *V*, attention is calculated as:

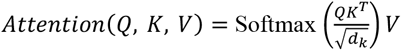

Here, *QK*^T^ computes similarity scores between the query and keys, scaled by 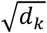 (the dimension of the keys) for numerical stability. The softmax function normalizes these scores into attention weights, which are then used to compute a weighted sum of the values *V*. This process allows the model to prioritize relevant information dynamically for each step in the output generation.

In our model, the interaction is performed bi-directionally using shared weight matrices (*W*_*Q*_, *W*_*K*_, *W*_*V*_) for the linear projections of the query, key and value inputs. First, the input encoding of the mRNA slice (*X* _*RNA*_) is projected to form the query (*Q* _*RNA*_ = *X* _*RNA*_*W*_*Q*_). Simultaneously, the siRNA encoding (*X*_*siRNA*_) is projected to form the key (*K*_*siRNA*_ = *X*_*siRNA*_*W*_*K*_) and value (*V*_*siRNA*_ = *X*_*siRNA*_*W*_*V*_). For each nucleotide in the mRNA, the attention mechanism scans the entire siRNA, resulting in a contextualized representation of mRNA (*H* _*RNA*_) informed by the most salient features of its binding partner.

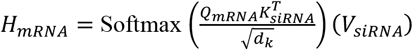

Then, the roles are reversed, but the same shared projection weights (*W*_*Q*_, *W*_*K*_, *W*_*V*_) are utilized. The siRNA sequence forms the query (*Q*_*siRNA*_ = *X*_*siRNA*_*W*_*Q*_), while the mRNA sequence forms the key (*K* _*RNA*_ = *X* _*RNA*_*W*_*K*_) and value (*V* _*RNA*_ = *X* _*RNA*_*W*_*V*_). This creates a contextualized representation of the siRNA (*H*_*siRNA*_), with its features now structurally informed by the mRNA target region.

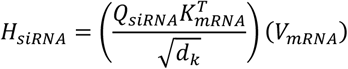

Finally, a single output vector is acquired by computing the element-wise average of the two attention output matrices.

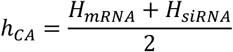

The shared bidirectional cross-attention mechanism captures the reciprocal nature of siRNA-mRNA interactions. It provides for the two sequences to be projected onto a common embedding space through the use of shared query, key and value projections for symmetrical information exchange. siRNA-mRNA pairing is not a one-way recognition process but a mutual alignment that is biologically meaningful. Each sequence refines its internal representation based on cues emanating from its partner through the bidirectional cross-attention mechanism, effectively mirroring the mutual recognition process inherent in RNA-RNA binding. This allows the model to learn higher-order relational dependencies and generalize to diverse sequence contexts.

#### 3.4.3. Thermodynamics Features and MLP

The thermodynamic stability profile of the siRNA duplex is one of the key determinants of its silencing efficiency. The energy asymmetry between the two ends of the duplex determines which strand is preferentially loaded into the RNA-induced silencing complex (RISC), thereby affecting the overall gene-silencing outcome. To quantitatively assess this property, feature engineering is performed to compute multiple thermodynamic parameters like the Watson-Crick pair free energy (ΔG) between adjacent nucleotide pairs, the total duplex free energy and the thermodynamic asymmetry between the 5’ and 3’ ends [33]. These features which are derived from established biophysical and thermodynamic models represent the duplex’s energy landscape and its contribution to efficacy prediction.

To perform the efficacy prediction, the feature representations obtained from the cross-attention module and SNN are concatenated with the pre-computed thermodynamic feature vector (*h*_*SNN*_). This gives a unified feature vector (*h*_*concat*_) that has sequential, structural and biophysical information:

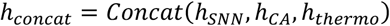

The comprehensive and high-dimensional feature vector is then passed into a MLP (Multi-layer perceptron) which acts as a regression head [34]. The MLP captures complex, non-linear relationships within the integrated features. Its output layer then transforms the learned representation into a single continuous value, representing the model’s predicted siRNA-mRNA efficacy score.

### 3.5. Loss Functions

The model is trained using a hybrid objective that integrates a regression loss for efficacy prediction and contrastive loss to align the latent representations of the siRNA and mRNA embeddings.

The primary objective is the Mean Squared Error (MSE) [6](PMID: 16025102) between the predicted and experimentally measured efficacies to perform regression:

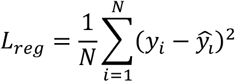

To encourage the model to differentiate effective from ineffective interactions, contrastive loss [35] between the siRNA and mRNA embeddings is obtained from the shared embedding space:

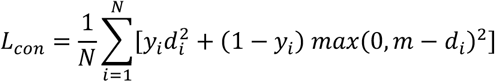

where *d*_*i*_ = ‖ *z* _*m,i*_ − *z*_*s,i*_ ‖_2_ denotes the Euclidean distance between the paired embeddings, *y*_*i*_ is a binary label derived from efficacy (1 for efficacy > 0.7, 0 otherwise [36], and *m* is contrastive margin.

The final loss is a weighted sum of the two objectives:

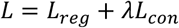

where *λ* controls the contribution of the contrastive term.

The joint optimization encourages the network to produce both accurate efficacy predictions and semantically align the siRNA-mRNA embeddings, improving generalization and interpretability.

### 3.6 Evaluation Metrics

The performance of the model is evaluated using both regression and classification-based metrics.

For regression, the metrics reported are:

Mean Squared Error (MSE) [10]: It is the mean squared difference between predicted and the experimentally validated efficacy values. It helps us understand how close the model’s predictions are to the truth values.

Pearson Correlation Coefficient (PCC): It is used to measure the strength and direction of the linear relationship between the predicted efficacy values and the actual efficacy values.

Spearman Correlation Coefficient (SPCC) [11]: It is used to measure the monotonic relationship between predicted values and ground truth values independent of scale.

For classification-based evaluation, siRNAs with efficacy > 0.7 are considered effective, and the rest are considered as ineffective. Using this binary threshold, Area Under Receiver Operating Characteristic Curve (AUC) is computed as a metric, showing the model’s ability to distinguish functional from non-functional siRNAs.

Using both regression and classification-based metrics is important as knockdown efficacy is inherently continuous, yet it is biologically interpreted in discrete terms such as effective or ineffective. This combined evaluation captures both the quantitative prediction accuracy and the functional relevance of the model in the biological context of RNAi.

## 4. Conclusion

The present work introduces SiaRNA, a deep learning framework for prediction of siRNA-mRNA efficacy, based on biologically aware feature engineering combined with a Siamese Neural Network and bidirectional cross-attention architecture. Our model leverages handcrafted sequence-derived features like nucleotide composition, GC content, base-pairing probabilities and nucleotide frequencies along with contextual sequence embedding. This allows the model to capture both intrinsic sequence properties and pairwise interaction dynamics. Encoding the siRNA and mRNA sequences into separate latent spaces before projecting them into a shared embedding space allows the model learn meaningful relational representations. The bidirectional cross-attention mechanism plays a major role in capturing mutual, context-dependent interactions which reflect the dynamic and reciprocal nature of RNA interference. The model is trained and validated on Dataset_HUVK and tested on the independent Simone dataset. SiaRNA yields an improvement in generalization and interpretability over previous approaches.

SiaRNA provides a dependable framework for the design of therapeutic siRNAs, allowing for the identification of more accurate siRNA candidates for gene silencing applications. Beyond therapeutics, the combination of feature-driven and attention-based modelling can enable broader gene regulation research and RNA-targeted interventions. This offers insights into the sequence-structure-functional relationships which regulate RNA-RNA interactions. This paves the way for integration with chemically modified siRNAs and the development of generalized RNA-targeted design approaches in diverse biological contexts.

## Author Contributions

Conceptualization, S.V., N.R., M.V.H., and N.V.R.; methodology, S.V., N.R., M.V.H., and N.V.R.; software, B.B.C.; validation, S.V., N.R., M.V.H., and N.V.R.; formal analysis, S.V., N.R., M.V.H., and N.V.R.; investigation, S.V., N.R., M.V.H., and N.V.R.; resources, B.B.C.; data curation, S.V., N.R., M.V.H., and N.V.R.; writing—original draft preparation, S.V., N.R., M.V.H., and N.V.R.; writing—review and editing, K.V.; visualization, B.B.C..; supervision, B.B.C., and K.V.; project administration, B.B.C.. All authors have read and agreed to the published version of the manuscript.

## Institutional Review Board Statement

Not applicable.

## Informed Consent Statement

Not applicable.

## Data Availability Statement

Data and code related to this study are available at https://github.com/drugparadigm/SiaRNA.

## Acknowledgements

We authors, S.V., N.R., M.V.H., N.V.R, B.B.C., and V.K, express our sincere gratitude to the Drugparadigm Research Lab for providing the necessary facilities and infrastructure that enabled the successful completion of this work.

## Abbreviations

The following abbreviations are used in this manuscript:

AGO2: Argonaute-2
AUC: Area Under Curve
CNN: Convolutional Neural Network
dsRNA: double-stranded Ribonucleic Acid
GNN: Graph Neural Network
GPU: Graphics Processing Unit
MLP: Multilayer Perceptron mRNA Messenger Ribonucleic Acid
MSE: Mean Squared Error
Nt: Nucleotide
PCC: Pearson Correlation Coefficient
RISC: RNA-Induced Silencing Complex
RNA: Ribonucleic Acid
RNAi: Ribonucleic Acid interference
RNase: Ribonuclease
ROC: Receiver Operating Characteristic
RPI: RNA-Protein Interaction shRNA short hairpin Ribonucleic Acid
siRNA: small-interfering Ribonucleic acid SNN Siamese Neural Network
SPCC: Spearman Correlation Coefficient SVD Singular Value Decomposition

